# An Early Olfactory Transcriptomic Signature of Tauopathy: Gbp2b Emerges as a Candidate Biomarker of Tau-Driven Neuroinflammation

**DOI:** 10.1101/2025.07.01.662324

**Authors:** Marion Dourte, Ayeh Bouloki, Marc Dieu, Malika Magomadov, Esther Paître, Nuria Suelves, Patricia Renard, Pascal Kienlen-Campard

**Author notes:** Contributed equally (Marion Dourte, Ayeh Bouloki).

## Abstract

Olfactory dysfunction is increasingly recognized as an early feature of neurodegenerative diseases such as Alzheimer’s disease. PS19 mice, a well-established tauopathy model, exhibit hallmarks of tau pathology—including hyperphosphorylated tau and pretangle formations—in various regions of the olfactory system. Notably, very recent data demonstrated that aberrantly hyperphosphorylated tau (pTau) was detected as early as 1.5 months of age in the olfactory epithelium (OE). This region contains olfactory sensory neurons projecting to the olfactory bulb (OB), where similar pTau pattern was also observed at this early stage. By 6 months, tau pretangles were evidenced in higher olfactory areas such as the piriform and entorhinal cortices. Given the early involvement of the OE and OB in tau pathology, we performed transcriptomic analyses at 3, 6, and 9 months to investigate the molecular pathways underlying tau pathology in these olfactory regions. Due to the OE’s peripheral location and anatomical accessibility, we also aimed in that respect to identify potential early biomarkers of tauopathy. The hippocampus, a key brain region affected in Alzheimer’s disease and related disorders, was included in the analysis as a comparative reference due to its known vulnerability and clinical relevance. Our analyses revealed region- and age-specific gene expression changes in PS19 mice. Functional enrichment analyses indicated a temporal progression of molecular alterations associated with tau pathology. We identified a subset of genes differentially expressed across different time points and/or regions. Among these, *Gbp2b* emerged as a particularly promising early biomarker candidate for tauopathy in the OE, showing consistent upregulation across tau pathological stages and brain regions.

**Graphical Abstract:** Created in BioRender. Kienlen-Campard, P. (2025) https://BioRender.com/k7xoypd

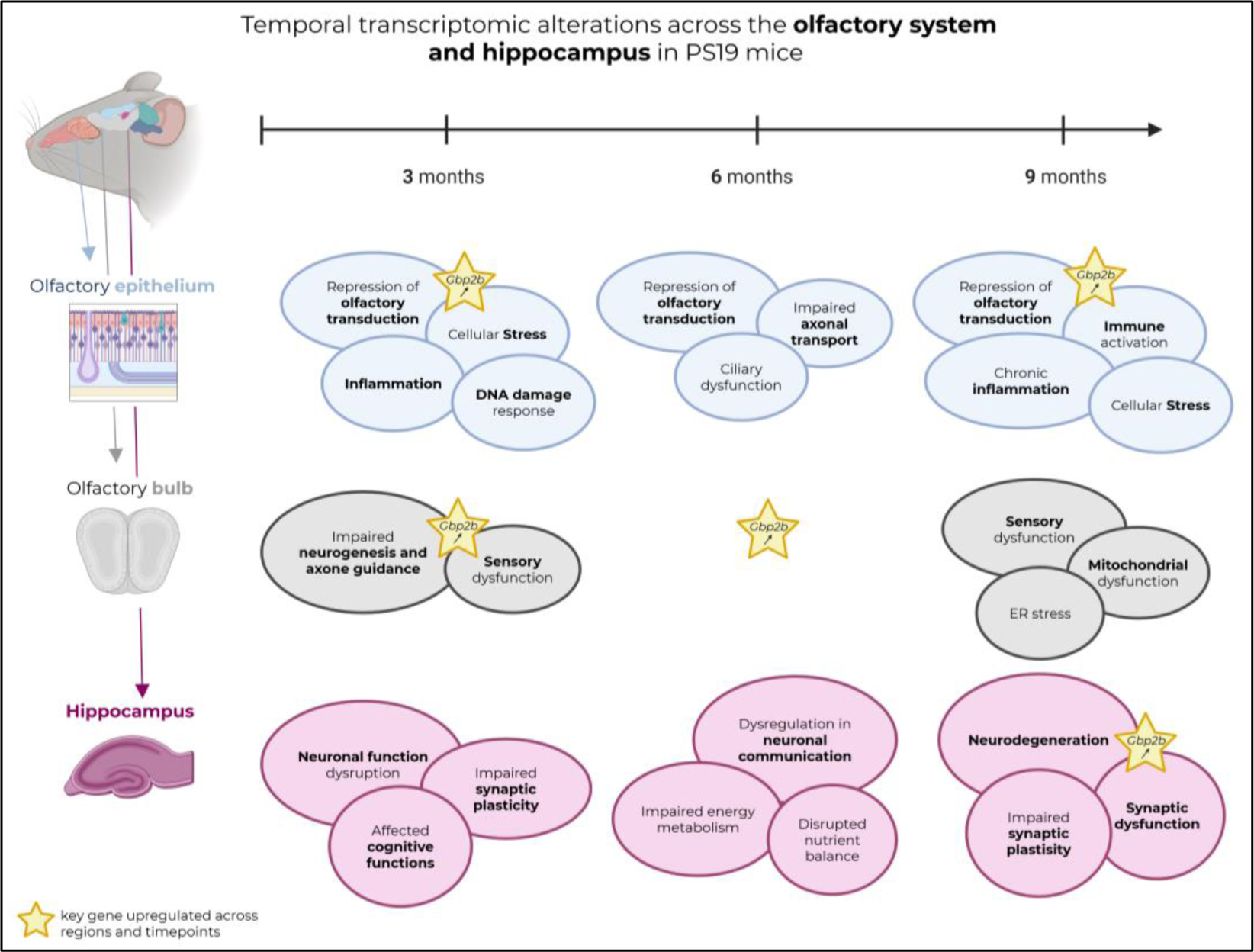

## Introduction

Alzheimer’s disease (AD) is defined by the presence of two histopathological lesions: extracellular amyloid-beta (Aβ) plaques, resulting from the aggregation of Aβ peptides, and intracellular neurofibrillary tangles (NFTs), composed of hyperphosphorylated tau protein [28]. Olfactory dysfunction is increasingly recognized as a feature of various neurodegenerative diseases (NDs). In AD, it affects approximately 90% of patients in the early stages and is characterized by decreased odor discrimination, increased olfactory threshold and olfactory memory loss [2, 3, 44]. It is thus considered a prodromal symptom of AD, often preceding cognitive impairment [18, 32, 40].

Transgenic mice that recapitulate major features of AD have been used to assess olfactory dysfunction and its underlying molecular mechanisms [18, 19, 40]. Particularly, PS19 mice are commonly employed to characterize tau pathology in the olfactory and central nervous systems [11]. These mice, also known as Tau P301S mice, were first described as a transgenic tauopathy mouse model by Yoshiyama *et al.* in 2007 [41]. They express mutant human microtubule-associated protein tau (MAPT) under the control of the mouse prion protein promoter (*Prnp*). The transgene encodes the 1N4R isoform harboring the disease-associated P301S mutation [41]. In this model, tau inclusions appear in key regions of the brain at 6 months and, by 9-12 months, extensive neurodegeneration and neuronal loss occur [41]. In our previous study, hyperphosphorylated tau protein (pTau, AT8-positive) was initially observed in olfactory regions such as the olfactory epithelium (OE) and olfactory bulb (OB) and later in other central nervous system (CNS) regions, including the piriform cortex, entorhinal cortex and hippocampus. We confirmed that pTau accumulates early in the OE of PS19 mice and subsequently spreads to the CNS following neuroanatomical pathways, including the olfactory and perforant pathways, ultimately reaching the hippocampus [11].

Given the early presence of tau pathology in the OE and the anatomical accessibility of this tissue, the OE represents a promising site for identifying potential biomarkers that could enable early diagnosis of tauopathies, including AD. In this study, we performed RNA sequencing in key regions of the olfactory and central nervous systems from PS19 and wild-type (WT) mice. We first examined differentially expressed genes (DEGs) in the OE, the primary sensory structure for smell. The OE, located in the nasal cavity, comprises olfactory sensory neurons (OSNs), supporting cells, and basal stem cells, which can generate OSNs throughout life [30]. Next, we investigated the OB, where OSNs project their axons, to form the outermost olfactory nerve layer. The OB consists of six layers where synapses occur between OSN axonal projections and other cells, such as mitral and tufted cells. Mitral cells, for instance, send projections to distant cortical regions, including the piriform and entorhinal cortices [4, 30]. Finally, we analyzed the hippocampus, a critical region for learning and memory that receives input from the entorhinal cortex [16]. Building on our previous study, in which we characterized tau pathology spread in these regions, we now aim to assess the early impact of this pathology on the olfactory system at the transcriptomic level.

While transcriptomic studies have been conducted on the cortex and hippocampus of PS19 mice [6, 15], no such studies have been performed on the OB or OE. Our study aims to fill this critical gap by applying RNA-sequencing analyses to these regions and evaluating the progression of molecular signatures associated with tauopathy over time.

## Methods

### Animals

PS19 mice (B6; C3-Tg(Prnp-MAPT*P301S)PS19Vle/J; Strain #008169), obtained from The Jackson Laboratory, were maintained on a C57BL/6 genetic background. Both wild-type (WT) and heterozygous PS19 mice were generated through controlled breeding. Males at 3, 6, and 9 months of age were used in this transcriptomic study. Females at 3 months were included for validation steps. The genotype of each animal was confirmed by PCR analysis of DNA isolated from ear or tail biopsies. Animals were housed in well-ventilated cages under a 12-hour light/dark cycle. They had ad libitum access to food and water throughout the study. Mice were individually identified via ear tags, and their health and behavior were closely monitored to ensure their well-being. All experimental protocols involving animals were approved in advance by the ethical committee at UCLouvain (2021/UCL/MD/018) and were carried out in strict accordance with institutional guidelines and national regulations for animal welfare. Humane endpoints were established to minimize any suffering, and appropriate steps were taken to ensure proper animal care throughout the duration of the study.

### Tissue Harvesting

All mice were euthanized by cervical dislocation to minimize stress and discomfort. Following euthanasia, the skin over the head was carefully removed to expose the cranium. The cranium was then opened, and the brain was dissected by making a sagittal cut along the midline, dividing the brain into two halves. From each half, the OB and the hippocampi were carefully extracted for further analysis. Subsequently, the mouse snout was dissected and cut along the midsagittal plane, separating the two nasal cavities. The OE was harvested from these cavities, identifiable by its distinctive yellow coloration, which is a hallmark of the tissue. All harvested tissues were immediately placed into pre-chilled isopentane, which was maintained at a freezing temperature to ensure rapid tissue preservation. The tissues were snap-frozen and subsequently stored at –80°C for long-term storage until further processing for analysis.

### Sample preparation for immunostaining

#### Decalcification

For immunostaining experiments, decalcification was needed to preserve the OE structure because of its localization within the nasal cavity [11]. The skin, eyes, and lower jaw were removed from the mouse head. The head was directly rinsed in phosphate-buffered saline (PBS) and subsequently fixed with modified Davidson’s fixative (mDF) (37% formalin, ethanol, glacial acetic acid, water) for four days under agitation at room temperature (RT). The heads were rinsed for 24 hours in PBS and transferred to OsteoRAL R (RAL Diagnostics, #320715-2500), a commercial decalcifier, for five days under agitation at RT. After decalcification, the mouse heads were embedded in paraffin.

#### Paraffin sections

Paraffin-embedded sections were obtained via a microtome. Coronal sections of 10 μm thickness were placed on Superfrost slides for immunohistochemistry.

### Immunohistochemistry

Paraffin-embedded sections of 10 μm were immersed in different solutions as follows: 2×5 minutes in toluene, 2×2 minutes in absolute isopropanol, and 30 minutes in 0,3% H_2_O_2_ in methanol. The slides were finally placed in distilled water. Then, the tissue sections on the slides were delimited using a DakoPen and 1X Tris-buffered saline (TBS) was placed on each paraffin section for 10 minutes. This washing step was repeated two times.

Then, the sections were blocked with 10% normal goat serum (NGS) diluted in TBS for one hour at room temperature. The primary antibody (AT8, 1/250, ThermoFisher, #MN1020) was prepared in 1% NGS, placed on each section, and left overnight at RT. The negative control sections were incubated with 1% NGS only.

The next day, the sections were rinsed three times for 10 minutes with TBS 1X. The secondary antibody (Anti-mouse biotin-labeled, 1/200, PerkinElmer, #NEF823001) was diluted with TBS 1X and incubated with the corresponding sections for 30 minutes at room temperature. This step was followed by 3 TBS rinses. Stabilized Elite ABC reagent (Vector Laboratories, #PK-7100) was added to each section for 30 minutes. After another washing step, ImmPACT® DAB EqV Substrate (Vector Laboratories, #VEC.SK-4103-100) was prepared. Equivalent amounts of reagent 1 and reagent 2 were mixed and incubated in each section. The reaction was stopped after three minutes by transferring the slides into distilled water. The sections were dehydrated by immersion in different solutions as follows: 2 minutes in 70% isopropanol, 2 minutes in 90% isopropanol, 2×2 minutes in absolute isopropanol and 3×2 minutes in toluene. The sections were mounted with DPX mounting medium (Sigma-Aldrich, #1.00579.0500) and left to dry overnight at room temperature. The images were captured via an Axioskop 40 microscope (Zeiss).

### Sample Preparation for RNA Sequencing

Tissues from the OE (pooled bilaterally), one OB, and one hippocampus were collected from PS19 and WT males at 3, 6, and 9 months (n=5 per tissue/genotype/time point; total n=90). RNA was extracted using TriPure™ Isolation Reagent (Roche) followed by chloroform phase separation. Further purification was performed with the ReliaPrep™ RNA Tissue Miniprep System (Promega), and RNA was resuspended in RNase-free water. RNA concentration and purity were assessed by measuring absorbance at 260 nm using a BioSpec-nano spectrophotometer. A260/A280 and A260/A230 ratios were used to evaluate sample purity (Shimadzu). RNA integrity was confirmed by RNA Quality Number (RQN) with a cutoff ≥8.0. Samples (800 ng RNA each) were sent to the GIGA Institute (Liège, Belgium) for RNA sequencing.

### RNA-sequencing Data Processing and Analysis

Raw Illumina reads underwent quality control with Fastp (v0.24.0), FastQC (v0.12.1), and MultiQC (v1.25.2) for adapter trimming and filtering of low-quality bases (>5% Ns). Clean reads were aligned to the GENCODE M36 (GRCm39) reference genome using STAR. Alignments were sorted with SAMtools (v1.21), and gene-level counts were generated with HTSeq (v2.0.5). Multidimensional scaling (MDS) plots using DESeq2-normalized counts assessed sample clustering. Differential expression analysis was conducted with DESeq2, applying thresholds of adjusted p-value <0.05 and |log2 fold-change| >0.58 to identify significantly DEGs.

### Functional Annotation and Enrichment

Gene Ontology (GO) and KEGG pathway enrichment analyses were performed using ClusterProfiler and g:Profiler in R (v4.2.1). Enriched terms (adjusted p < 0.05) were categorized into biological process, molecular function, and cellular component domains, providing insight into the molecular pathways disrupted in tauopathy. Results were visualized via bar plots, dot plots, and network diagrams to facilitate interpretation.

### qRT-PCR Validation of Candidate Biomarkers

To experimentally validate candidate biomarkers, we performed quantitative reverse transcription PCR (qRT-PCR) on OE samples. Tissue was collected from 3, 6, 9-month-old male and female PS19 transgenic mice and age-matched wild-type mice (n = 4 or 5/group).

Total RNA was extracted using TriPure™ Isolation Reagent (Roche) followed by chloroform phase separation. Further purification was performed with the ReliaPrep™ RNA Tissue Miniprep System (Promega), and RNA was resuspended in RNase-free water. Concentration and purity were assessed via absorbance at 260 nm using a BioSpec-nano spectrophotometer (Shimadzu). cDNA was synthesized using iScript cDNA synthesis Kit (Bio-Rad). qRT-PCR was performed using SYBR Green Master Mix on a Bio-Rad CFX96 real-time PCR system. Each reaction was run in technical triplicates. Gene expression levels were normalized to housekeeping gene (Table S6)(Gapdh), and relative expression was calculated using the 2^−ΔΔCt^ method. Data were first tested for normality using the Shapiro–Wilk test. When data followed a normal distribution, parametric statistical analysis was performed using a two-tailed unpaired Student’s *t*-test. For data that did not meet normality assumptions, non-parametric analysis was conducted using the Mann–Whitney *U* test. A *p*-value < 0.05 was considered statistically significant. Candidate genes were selected for validation based on magnitude of differential expression, overlapping genes and OE-specificity.

### Protein Sample Preparation

Samples preparation and analysis were performed in the Mass Spectrometry Facility from the University of Namur. At 6 months, OE (pooled), one OB, and one hippocampus were collected from PS19 and sex-matched WT controls mice (n=10 per group; total n=30). Tissue samples (∼10 mg) were homogenized in 200 µL urea lysis buffer (8 M urea, 2% SDS, 1 mM EDTA, 50 mM Tris-HCl, pH 8.0) supplemented with protease and phosphatase inhibitors. Homogenization employed zirconium beads and a FastPrep-24 (MP Biomedicals) device (3 × 20 s cycles at 6.5 m/s with 2-min cooling intervals at 4°C). Samples were sonicated using a Bioruptor (Diagenode)(5 × 30 s bursts, 60 s intervals on ice), then centrifuged at 14,000 rpm for 5 min at 4°C. The clarified supernatant was collected, and protein concentration determined by Pierce 660 nm assay (ThermoFisher). Protein aliquots were stored at −80°C prior to proteomic analysis.

### Protein Digestion

The samples were treated using Filter-aided sample preparation (FASP) using the following protocol. To first wash the filters, 100 µl of 1% formic acid was added to each Microcon 30 filter unit (Millipore), followed by centrifugation at 14500 rpm for 15 minutes. For each sample, 40 µg of protein was brought to a final volume of 150 µl using 8M urea buffer (urea 8 M in buffer Tris 0.1 M at pH 8,5) and was loaded into a column and centrifuged at 14500 rpm for 15 minutes. The filtrate was discarded and the columns were washed three times by adding 200 µl of urea buffer followed by a centrifugation at 14500 rpm for 15 minutes. For the reduction step, 100 µl of 8mM dithiothreitol (DTT) was added to the columns and mixed for 1 minute at 400 rpm with a thermomixer before an incubation of 15 minutes at 24 °C. Samples were then centrifuged at 14500 rpm for 15 minutes, the filtrate was discarded and the filter was rinsed by adding 100 µl of urea buffer, followed by another centrifugation at 14500 rpm for 15 minutes. An alkylation step was performed by adding 100 µl of 50mM iodoacetamide (IAA), in urea buffer) to each column, followed by mixing at 400 rpm for 1 minute in the dark. The samples were then incubated for 20 minutes in the dark and centrifuged at 14500 rpm for 10 minutes. To remove the excess of IAA, 100 µl of urea buffer was added and the samples were centrifugated at 14500 rpm for 15 minutes. To quench residual IAA, 100 µl of DTT was placed on the column, mixed for 1 minute at 400 rpm and incubated for 15 minutes at 24 °C before centrifugation at 14500 rpm for 10 minutes. To remove the excess of DTT, 100 µl of urea buffer was added to the column and centrifuged at 14500 rpm for 15 minutes. The filtrate was discarded, and the column was washed three times by adding 100 µl of sodium bicarbonate buffer 50 mM (ABC) in ultrapure water, followed by a centrifugation at 14500 rpm for 10 minutes. The last 100 µl of buffer was left at the bottom of the filter unit to avoid any evaporation in the column. The protein digestion process was performed by adding 80 µl of mass spectrometry-grade trypsin (1/50 in ABC buffer) to the column, followed by mixing at 400 rpm for 1 minute before an incubation overnight at 24°C in a water saturated environment. The Microcon columns were placed into LoBind tubes of 1.5 ml and centrifuged at 14500 rpm for 10 minutes. 40 µl of ABC buffer was placed on the column before centrifugation at 14500 rpm for 10 minutes. 10 % Trifluoroacetic acid (TFA) in ultrapure water was added to the content of the LoBind Tube to obtain a final concentration of 0.2 % TFA. The samples were dried in a SpeedVac to a final volume of 20 µl and transferred into injection vials.

### DIA-MS Protein Data Processing and Analysis

DIA-MS Protein Data Processing and Analysis was used as target validation. The protein digest was analyzed using nano-LC-ESI-MS/MS time TOF HT (Bruker, Billerica, MA, USA) coupled with an UHPLC nanoElute 2 (Bruker).

Peptides were separated using a nanoUHPLC system (nanoElute2, Bruker) equipped with a 150 μm inner diameter × 25 cm column packed with 1.5 μm C18 particles (PepSep, Bruker). A CaptiveSpray emitter with a preassembled union was used to interface the column with the CaptiveSpray ion source (Bruker). The liquid chromatography mobile phases consisted of solvent A: water with 0.1% formic acid (v/v), and solvent B: acetonitrile (ACN) with 0.1% formic acid (v/v). Samples were loaded directly on the analytical column at a constant pressure of 800 bar. The digest (1 µl) was injected, and the organic content of the mobile phase was increased linearly from 5% to 27 % B in 56 minutes at a flow rate of 600 nl/min, from 27 % to 35% B in 7 minutes, from 35% to 95 % B in 1 minute. Data acquisition on the tims TOF HT was performed using Hystar 6.3. Mass-spectrometric analyses were carried out using the data-independent parallel accumulation serial fragmentation acquisition method (DIA-PASEF) [22]. The DIA-PASEF acquisition scheme was 1 MS1 ramp and 6 MS2 ramps with 18 windows. Mass range was restricted from 300 to 1200 m/z and ion mobility was set from 0.6 to 1.4 1/K_0_.

DIA-NN 2.0 [10] was used to process the raw data with an in silico spectral library of 54644 proteins and 22086 genes created from Mus Musculus proteome database (UNIPROT) and a contaminants database.

Trypsin/P was used with a maximum of 1 missed cleavage. Variable modifications on peptides were set to methionine oxidation, and fixed modification was set to carbamidomethylation on cysteine. The maximum number of variable modifications on a peptide was set to 3. The peptide length for the search ranged from 7 to 30 amino acids. m/z ranges were specified as 300 to 1400 for precursors and 100 to 1700 for fragment ions. Both MS1 and MS2 mass accuracy were set to 15ppm. Scoring was set to generic. Match-between-run (MBR) and protein inference options were checked. RT-dependent cross-run normalization and QuantUMS (high precision) options were selected for quantification. A 1% FDR cutoff was applied at the precursor level.

In order to perform the differential expression analysis, protein group matrix from DIA-NN was loaded to the web application DIA-Analyst from Monash University [43]. Adjusted p-value cutoff was set to 0.05 and log2 fold changed cutoff to 1. No imputation and t-statistics-based FDR correction were applied.

## Results

### Early Detection of Hyperphosphorylated Tau in the Olfactory Epithelium and Olfactory Bulb of PS19 Mice

As previously described [11], the presence of hyperphosphorylated tau (pTau) was detected using the AT8 antibody (pTau Ser202, Thr205) in coronal sections of the OE from WT and PS19 mice. To preserve OE integrity, a specific decalcification protocol was established. AT8 immunoreactivity was observed exclusively in PS19 mice, localized to OE olfactory sensory neurons (OSNs) and axon bundles (ABs) within the underlying lamina propria (Fig. 1a-c) at 3 months of age.

**Fig. 1.**
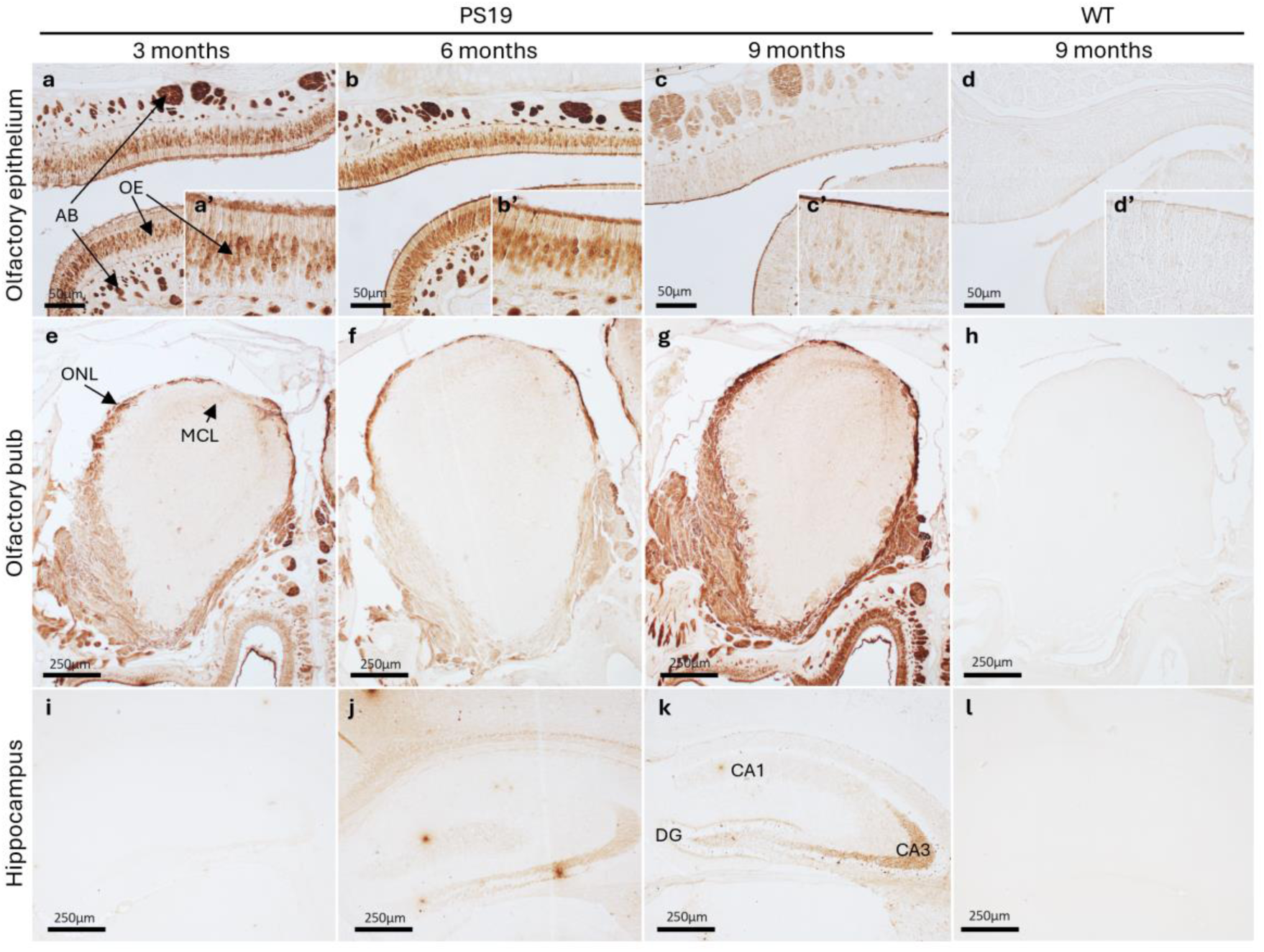
pTau expression in the OE, OB, and hippocampus of PS19 mice. The OE, OB and hippocampus from 3-month (a, e, i), 6-month (b, f, j), 9-month-old PS19 mice (c, g, k), and 9-month-old WT mice (d, h, l) were fixed and stained with the **AT8** antibody (pTau (Ser202, Thr205)). pTau recognized by the **AT8** antibody is found in the OSNs of the OE and in the AB located in the lamina propria at 3 months of age in PS19 mice (a). pTau is also found in the ONL and MCL of the OB from PS19 mice at 3 months of age (e). pTau is detected in the DG and CA3 of the hippocampus from PS19 mice from 6 months of age (j, k). All the mice used were males. High-magnification views of the OE are shown in the insets (a’, b’, c’, d’). ABs: axon bundles, OE: olfactory epithelium, ONL: olfactory nerve layer, MCL: mitral cell layer, DG: dentate gyrus, CA: Cornu Ammonis

In the OB of PS19 mice, pTau was also detected in the olfactory nerve layer (ONL) – formed by the OSNs projections – and mitral cell layer (MCL) at 3 months of age (Fig. 1e-g). AT8 signal emerged in the dentate gyrus (DG) and CA3 hippocampal subregions at 6 months (Fig. 1j), with increased intensity observed at 9 months (Fig. 1k). No pTau signal was detected in the OE, OB, or hippocampus of 9-month-old WT mice (Fig. 1d, h, l), confirming the specificity of AT8 staining to PS19 mice.

The early detection of pathological tau in key olfactory system regions, including the OE and OB, prompted us to perform transcriptomic analyses in these areas to uncover molecular alterations underlying tau pathology at different stages of the pathology.

### Transcriptomic Profiling Reveals Age- and Region-Specific Gene Expression in PS19 Mice

To characterize tauopathy progression, RNA sequencing was performed on the hippocampus, OB, and OE of PS19 and WT mice at 3, 6, and 9 months. Heatmaps, PCA and Euclidean distance analyses confirmed robust separation by genotype and age across all regions (Fig. 2, Fig. S1).

**Fig. 2.**
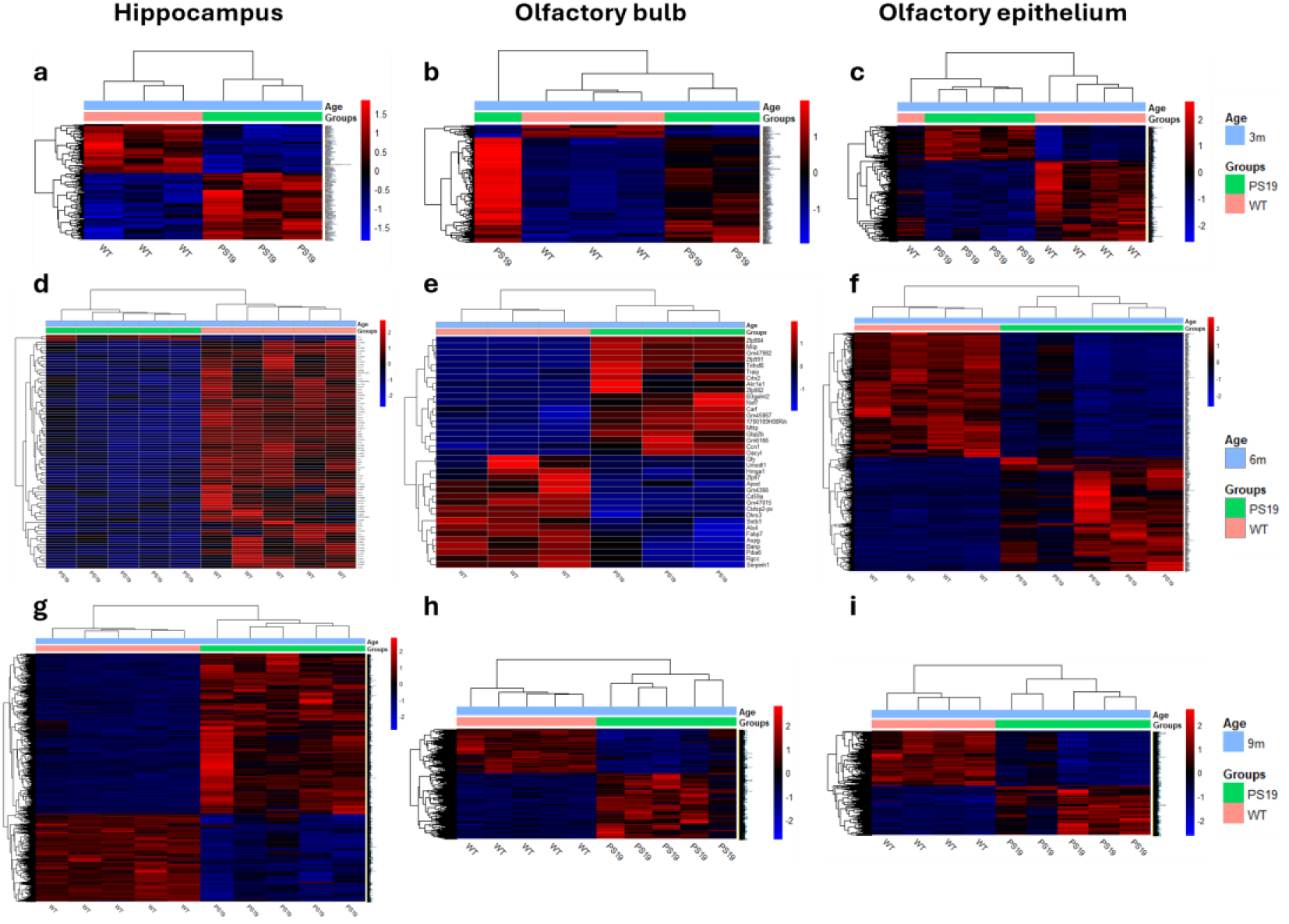
Hierarchical clustering heatmaps of normalized gene expression levels across multiple brain and olfactory regions in PS19 and WT mice at 3, 6, and 9 months. Heatmaps are shown for the hippocampus (a, d, g), OB (b, e, h), and OE (c, f, i). Gene expression was assessed over time at 3 months (a–c), 6 months (d–f), and 9 months (g–i). Rows represent differentially expressed genes, and columns correspond to individual samples (PS19 in green; WT in pink). Color scale indicates relative expression levels, with blue representing low expression and red representing high expression. N = 3-5 mice per group.

At 3 months, the OE exhibited the most transcriptomic disruption (hippocampus: 160, OB: 239, OE: 1023 DEGs). DEG numbers decreased in the hippocampus and OB by 6 months but remained high in the OE (1031), suggesting sustained dysfunction in the olfactory system. At 9 months, DEGs sharply increased across all regions (Hippocampus: 4599, OB: 2896, OE: 3665), aligning with disease progression. The top DEGs per anatomical region and time point are listed in **Table 1**.

**Table 1.**
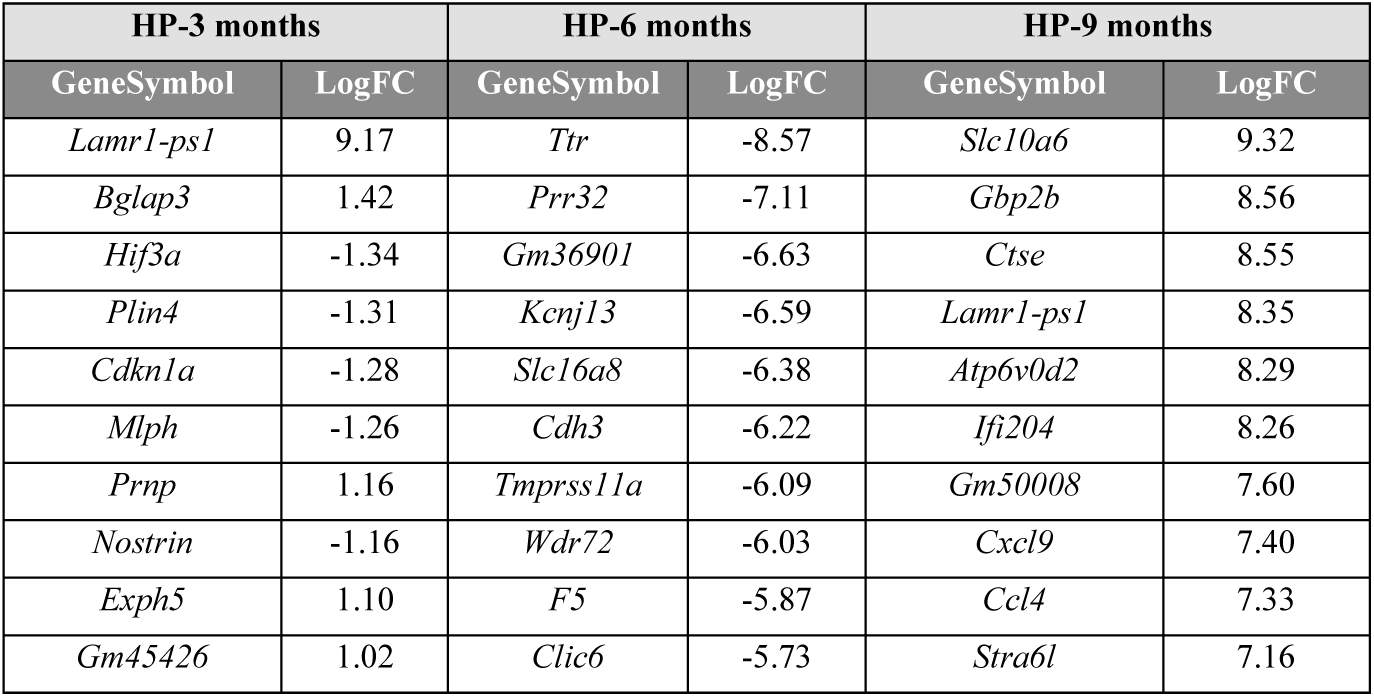
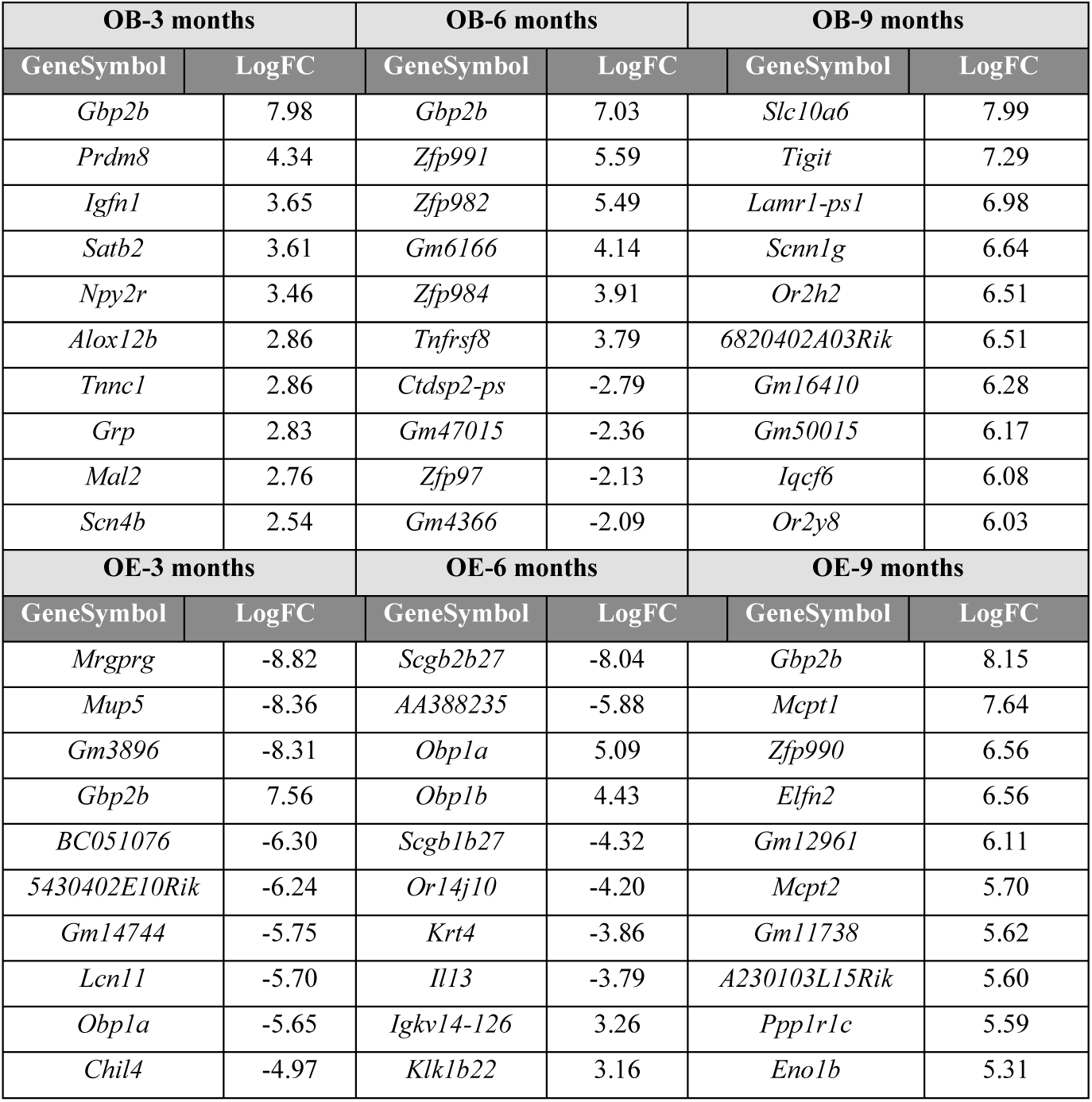
Top 10 DEGs ranked by the absolute logFC values across three regions (hippocampus, OB, and OE) at three time points (3, 6, and 9 months) in PS19 transgenic mice.

### Functional Enrichment Analysis Highlights Temporal Progression of Tauopathy

GO and KEGG pathway analysis of DEGs showed age- and region-specific enrichment. In the hippocampus, early changes (3 months) involved processes related to cognition, extracellular matrix organization, and cAMP signaling (Fig. 3a). At 6 months, signaling and metabolic disruptions emerged (Fig. 3b), while by 9 months, enrichment shifted toward neurodegeneration, synaptic dysfunction, and circadian regulation (Fig. 3c). In the OB, early DEGs were linked to axon guidance and neuroactive signaling (Fig. 4a). At 6 months, enrichment was minimal, suggesting transient stabilization. By 9 months, DEGs in the OB showed strong enrichment in olfactory signaling, endoplasmic reticulum stress, and synaptic targeting (Fig. 4b). The OE exhibited early signs of stress and immune activation at 3 months, with enrichments in sensory perception, DNA damage response, and p53 signaling (Fig. 5a). By 6 and 9 months, structural (cilium organization), inflammatory (TNF, NF-κB), and metabolic pathways (HIF-1) were increasingly dysregulated (Fig. 5b, c). Repression of olfactory transduction pathways was consistent with declining olfactory function. These findings highlight the progressive features of tauopathy and suggest that cellular stress, inflammation, and disrupted synaptic and neuronal functions play a key role in tau-related neurodegeneration across the studied regions. The olfactory system appears to be an early and vulnerable target of tau accumulation, with significant implications for sensory and cognitive functions in tauopathies.

**Fig. 3.**
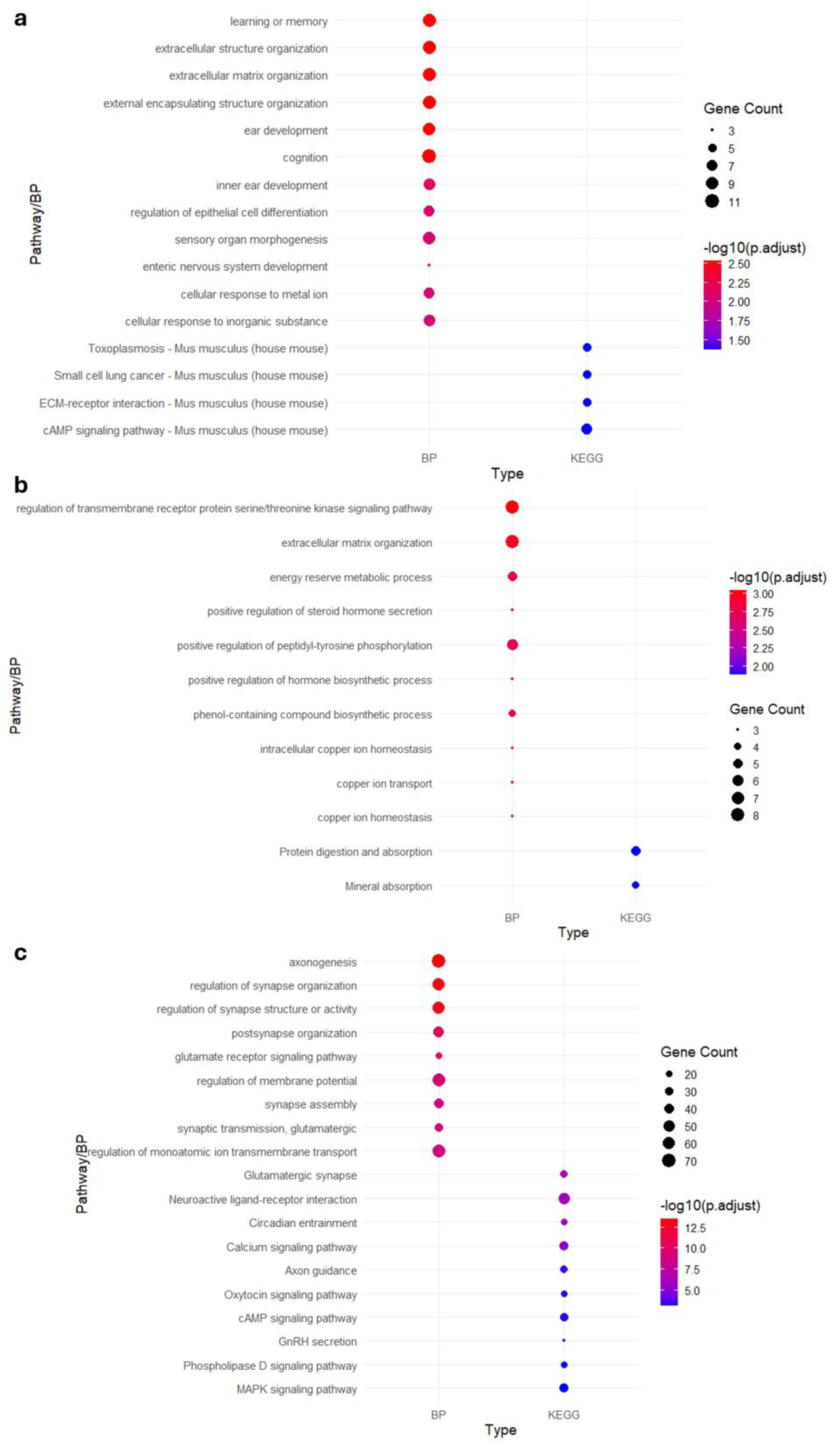
Enrichment analysis of DEGs in the hippocampus at 3 (a), 6 (b), and 9 (c) months. Dot plot showing significantly enriched terms among DEGs in these conditions. The y-axis lists enriched terms from GO Biological Processes (BP) and KEGG pathways, as indicated on the x-axis. Dot size represents the number of DEGs associated with each term (Gene Count), while dot color reflects statistical significance, shown as –log₁₀(adjusted p-value).

**Fig. 4.**
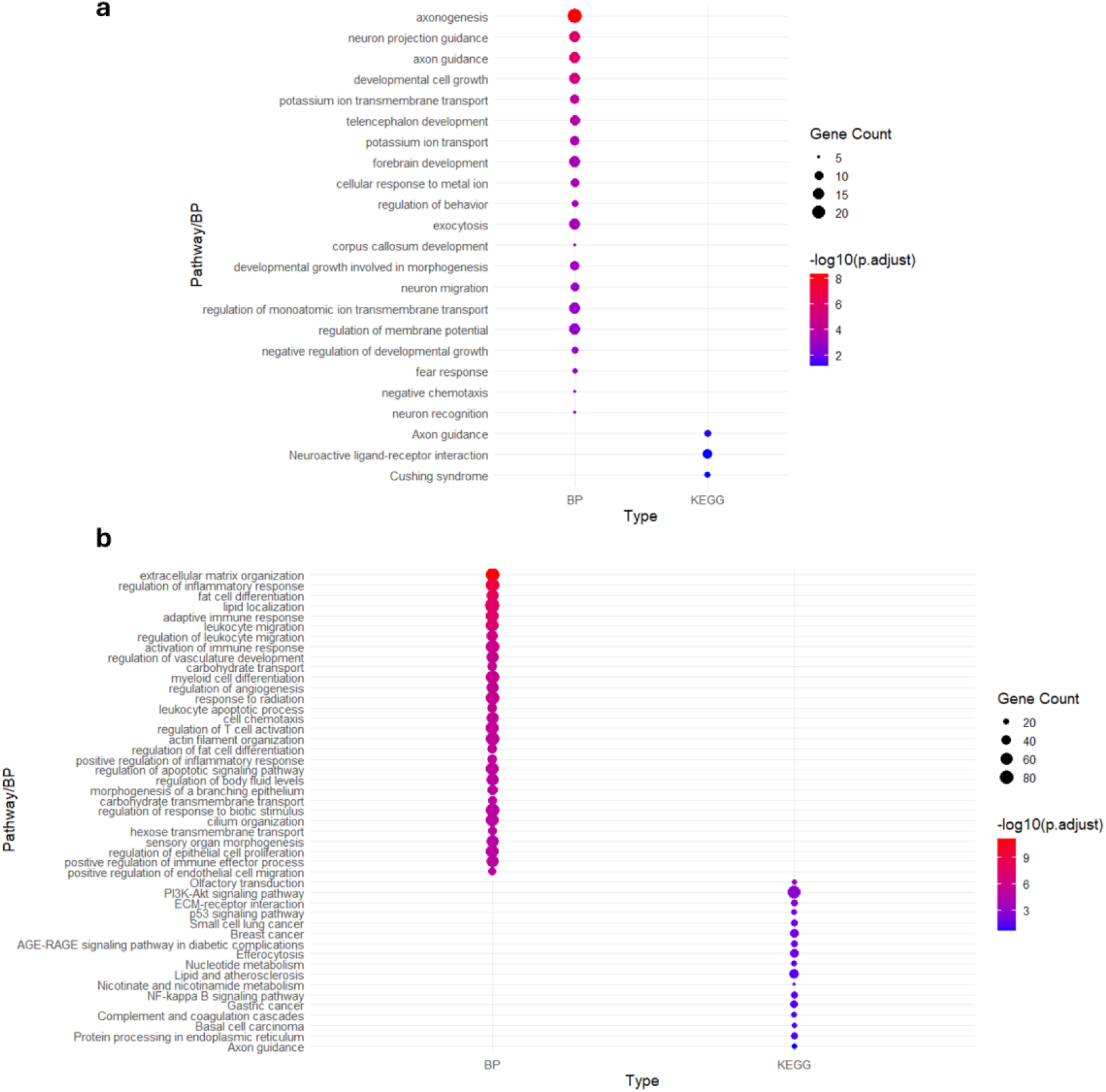
Enrichment analysis of DEGs in the OB at 3 (a) and 9 (b) months. No significantly enriched pathways in the OB at 6 months. Dot plot showing significantly enriched terms among DEGs in these conditions. The y-axis lists enriched terms from GO Biological Processes (BP) and KEGG pathways, as indicated on the x-axis. Dot size represents the number of DEGs associated with each term (Gene Count), while dot color reflects statistical significance, shown as –log₁₀(adjusted p-value).

**Fig. 5.**
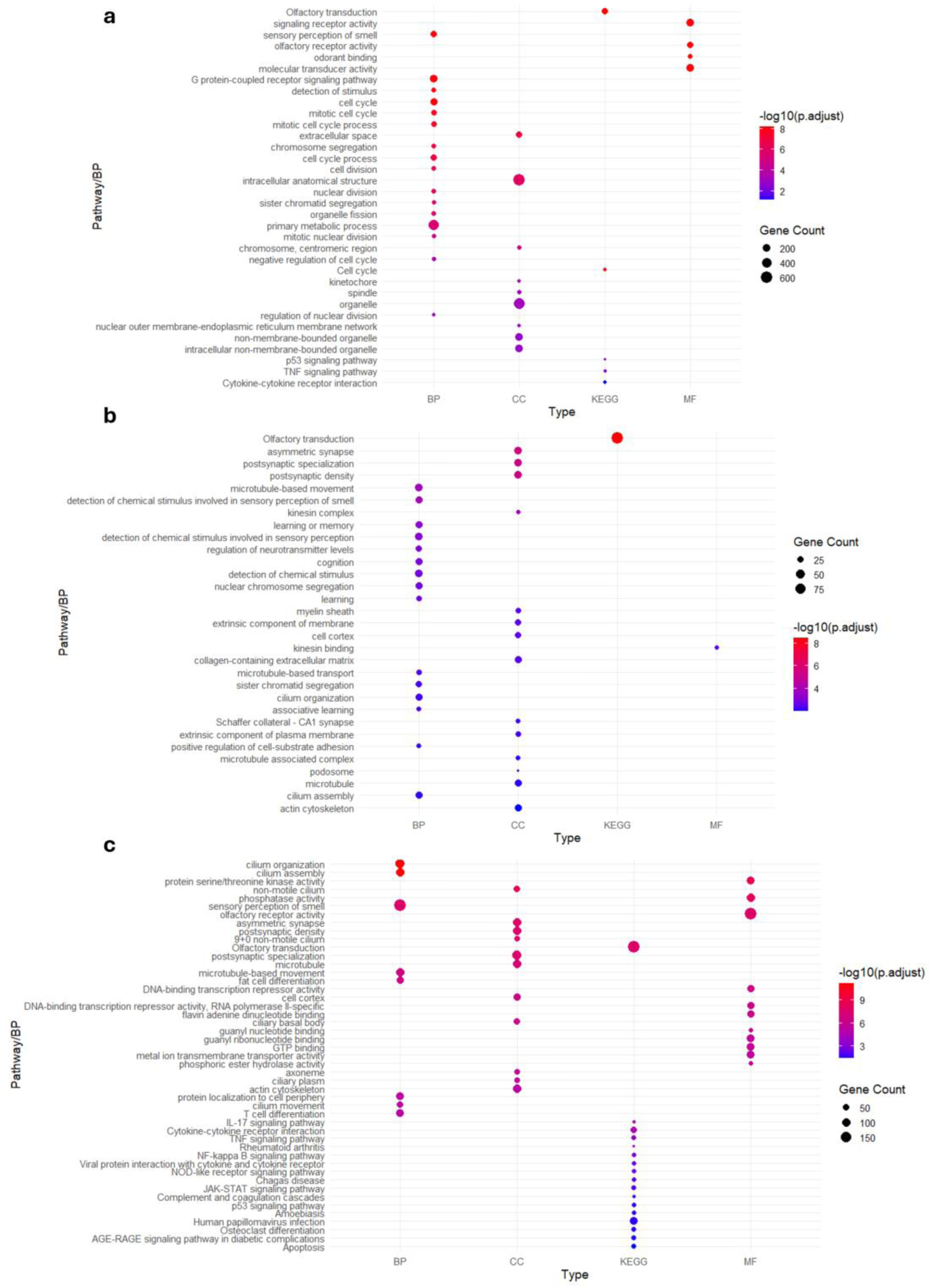
Enrichment analysis of DEGs in the OE at 3 (a), 6 (b), and 9 (c) months. Dot plot showing significantly enriched terms among DEGs in these conditions. The y-axis lists enriched terms from GO Biological Processes (BP), Cellular Component (CC), KEGG pathways, and Molecular Function (MF), as indicated on the x-axis. Dot size represents the number of DEGs associated with each term (Gene Count), while dot color reflects statistical significance, shown as –log₁₀(adjusted p-value).

### Spatiotemporal Overlap of Differentially Expressed Genes Reveals Region- and Age-Specific Responses to Tau Pathology

To investigate the spatiotemporal dynamics of gene expression in tauopathy, we examined the overlap of DEGs across the hippocampus, OB, and OE in PS19 mice at 3, 6, and 9 months of age. Venn diagrams and DEG comparisons were used to identify common and region-specific transcriptional responses during pathology progression.

At 3 months, overlap analysis revealed two DEGs—*Wnt9a and Prnp*—shared across all three regions (Fig. 6a). *Prnp* was consistently upregulated in all studied regions and may represent a neuronal stress response linked to tau accumulation. *Wnt9a* showed region-specific regulation, being upregulated in the hippocampus and OB but downregulated in OE. Given Wnt signaling’s role in neurogenesis and plasticity [25], this pattern may reflect compensatory responses in hippocampus/OB and impaired regenerative capacity in OE, potentially contributing to early olfactory dysfunction—a hallmark of Alzheimer’s and Parkinson’s disease [9, 20]. Further overlap between OB and OE identified 19 shared DEGs. Genes such as *Fosl2* and *Nr4a3* were downregulated in both regions, while others (e.g., *Gramd2a, Cgref1*) displayed opposing expression, indicating region-specific metabolic or stress responses. Immune-related genes, including *Gbp2b*, and Wnt pathway regulators *Dkk3* and *Wnt4*, were upregulated in both regions, suggesting activation of inflammatory and neurogenic pathways. Shared DEGs between hippocampus and OE (n = 13) included genes with both uniform and opposing expression patterns. For instance, *Ldlr* and *Mansc1* were upregulated in hippocampus but downregulated in OE, suggesting differential vulnerability to lipid dysregulation and stress. Uniformly downregulated genes—*Tsc22d3, Recql4, and Nkd2*, among others—highlighted common impairments in DNA repair, stress resilience, and synaptic function. Upregulated genes such as *Gm56721* further underscore overlapping responses to tau burden.

**Fig. 6.**
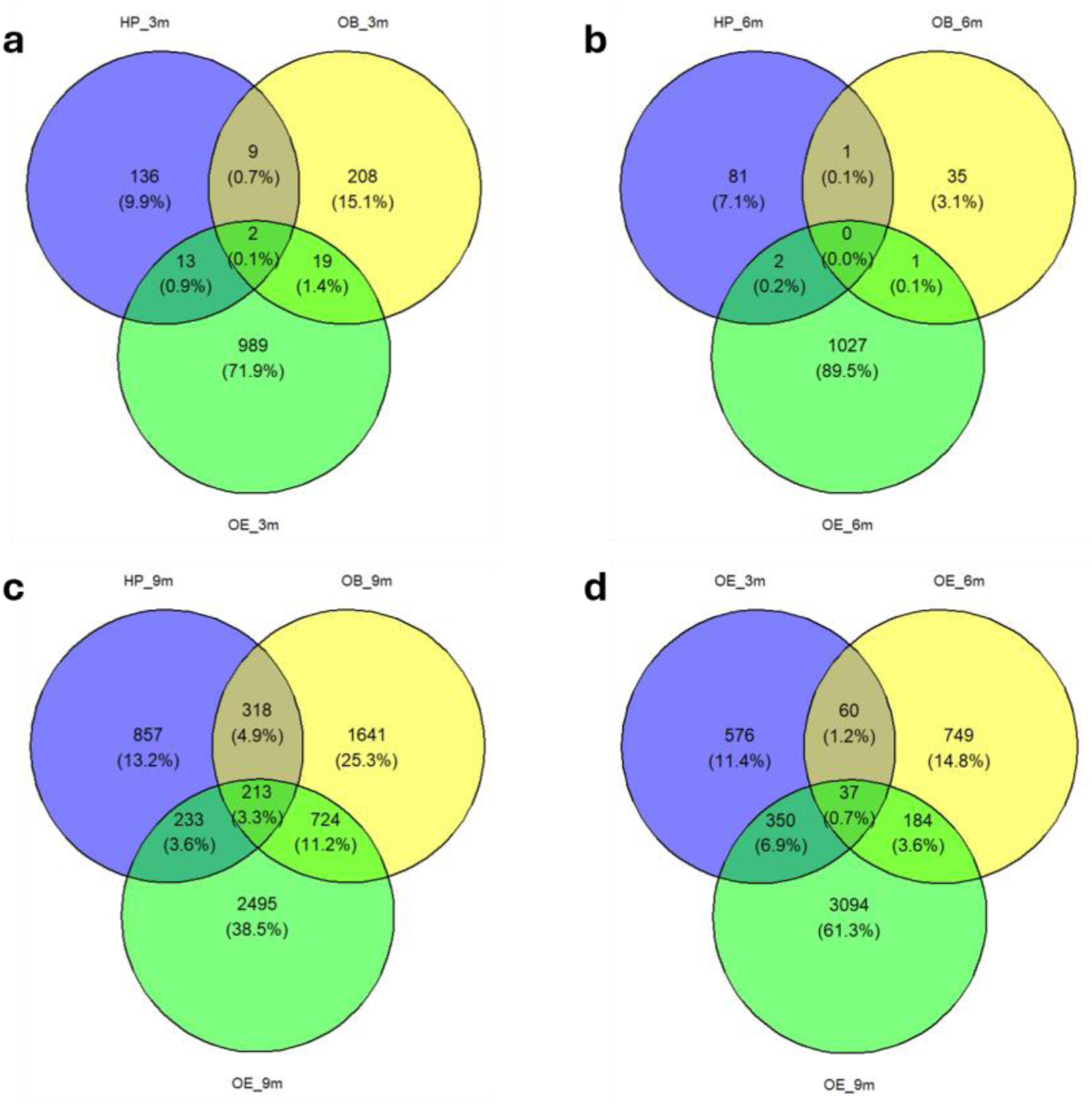
Venn diagrams showing the overlap of DEGs across regions and time points in PS19 mice. Overlap of DEGs among the hippocampus, OB, and OE at 3 months (a), 6 months (b), and 9 months (c). The majority of DEGs are specific to the OE, with limited overlap across regions at early time points and greater overlap observed at 9 months. Comparison of DEGs in the OE across 3, 6 and 9 months (d), showing progressive change over time. Percentages within the diagrams represent the proportion of total genes in each category.

At 6 months, the number of shared DEGs decreased, indicating increased regional specificity with aging (Fig. 6b). Only *Sulf1* and *Crb2* were common to hippocampus and OE, and *Gfy* to OB and OE. Notably, *Sulf1* and *Crb2* were upregulated in OE but downregulated in hippocampus, potentially reflecting continued regenerative activity in OE versus degenerative changes in hippocampus. *Gfy* upregulation in OB and downregulation in OE may indicate disrupted olfactory neurogenesis and connectivity.

At 9 months, DEG overlap increased substantially, with 213 genes shared across all three regions (3.3% of total DEGs), indicating a convergence of disease-associated molecular alterations (Fig. 6c). In hippocampus and OE, overlapping DEGs were enriched in synapse-related functions and ciliary components associated with long-term potentiation and calcium signaling—critical pathways for cognitive and sensory function. A large set of 724 overlapping DEGs (11.2%) between OB and OE included genes involved in DNA replication, cytokine signaling, apoptosis, and synapse pruning, emphasizing shared neurodegenerative mechanisms in olfactory regions.

Across all ages, multiple guanylate-binding proteins (GBPs) were consistently upregulated in OB and OE. These proteins are involved in immune responses, particularly in regulating inflammation, antiviral defense, and immune signaling, suggesting a sustained neuroinflammatory response in olfactory regions—a process increasingly linked to tauopathy progression [42].

Lastly, 37 genes in OE were differentially expressed across all time points in PS19 mice (Fig. 6d). 16 of these genes, including *Depdc1a, Sox4, and Hmgb2*, shifted from downregulation at 3 months to upregulation at 6 and 9 months. Many of these genes are involved in cell cycle regulation, signal transduction, and neurogenesis, which are key processes for maintaining the structural integrity of OSNs. This temporal pattern suggests an age-dependent compensatory response aimed at restoring homeostasis, particularly in pathways involving cell cycle regulation, neurogenesis, and DNA repair. In contrast, 21 genes were found to be upregulated at 3 months but downregulated at 6 and 9 months. Many of these genes encode olfactory receptors (ORs) such as *Or10g7*, *Or8b51*, or *Or5d39* suggesting an early overactivation of olfactory signaling, followed by a decline as disease progresses. This shift likely reflects impaired olfactory transduction and OSNs loss, aligning with the known early olfactory deficits in NDs like AD and Parkinson’s disease [9, 20].

Collectively, these findings reveal early and progressive transcriptomic changes in response to tau accumulation, with distinct and shared regional signatures. The OE, with its neurogenic capacity, displays dynamic regulatory shifts that are likely to contribute to early sensory impairments observed in tauopathies.

### Transcriptomic Signatures in the OE of PS19 mice as Early Biomarkers of Tau Pathology

A secondary objective of this study was to uncover early phenotypic alterations that may act as potential biomarkers for the early detection of tau-related NDs. The OE, due to its anatomical accessibility, serves as a valuable diagnostic tool compared to CNS regions like the OB or hippocampus, which are challenging to access in living organisms. While these regions are crucial for understanding the mechanisms driving tau pathology, our focus was on identifying potential biomarkers by analyzing DEGs in the OE. The 3-month time point was particularly relevant for early diagnosis, as these molecular changes emerge even before pathology develops in regions such as the hippocampus. We established a panel of candidate biomarkers (Table 2), from which a subset was selected for validation by qRT-PCR.

**Table 2.**
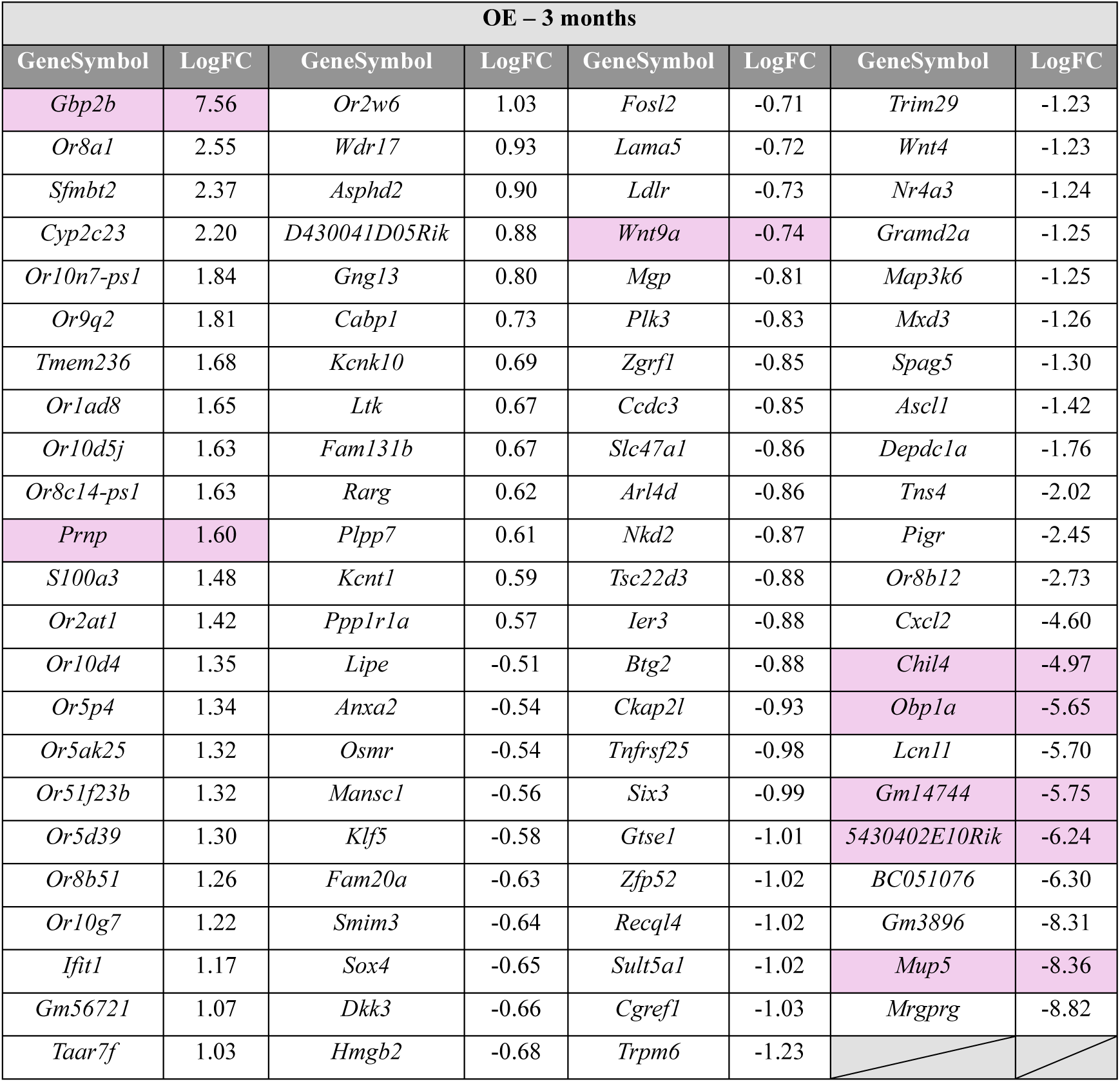
Biomarker panel in the OE at 3 months. Selection based on the DEGs in the OE at 3 months, and the overlapping genes analyses. The candidates validated by qRT-PCR are highlighted in pink.

We chose to perform the validation in a male cohort, consistent with the RNA-sequencing analysis, and additionally included a female cohort to assess potential sex-related differences in gene expression. We first validated *Prnp* increased expression in the OE and OB, since it was one of the two candidates upregulated across all three regions. The increase was significant from 6 months in the OE (Fig. 7a) and from 3 months in the OB (Fig. 7b).

**Fig. 7.**
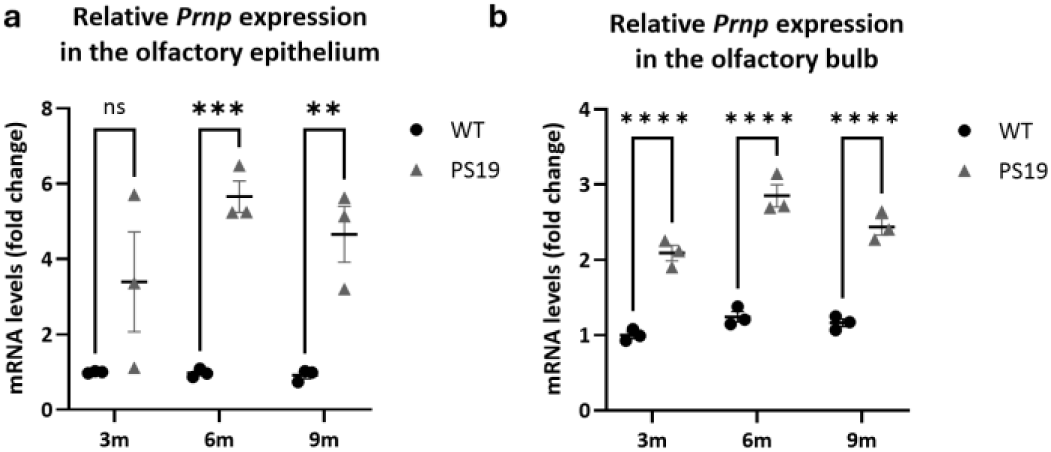
*Prnp* expression in the OE and OB of WT and PS19 mice. mRNA levels of *Prnp* expression were measured by RT-qPCR in OE and OB extracts from WT and PS19 mice at 3, 6, and 9 months. *Prnp* is significantly increased in the OE of 6- and 9-month-old PS19 mice (a). \*\**P* < 0.01, \*\*\**P* < 0.001. (Two-way ANOVA, n=3). *Prnp* is significantly increased in the OB of 3-, 6-, and 9-month-old PS19 mice (b). \*\*\*\**P* < 0.0001. (Two-way ANOVA, n=3).

Then we focused on several candidates that might be of interest for further validation among the top 10 DEGs in the OE at 3 months (*Mrgprg, Mup5, Gm3896, Gbp2b, BC051019, 5430402E10Rik, Gm14744, Lcn11, Obp1a, Chil4*)(Table 1). We further investigated other candidates that showed an almost perfect segregation in the box plots, with the formation of two clearly distinct groups (WT and PS19), such as *Mup5, Pon1,* and *Tsc22d1* (Fig. S2). Finally, we also checked for *Wnt9a* expression since it showed region-specific regulation: upregulated in hippocampus and OB but downregulated in OE.

qRT-PCR analysis confirmed the trend of reduced expression of *Mup5, 5430402E10Rik, Gm14744,* and *Obp1a* both in males and females, consistent with RNA-sequencing data (Fig. 8b-e). However, these differences did not reach statistical significance in all conditions, likely due to high inter-individual variability or low baseline expression. *Pon1* and *Tsc22d1* showed more distinct clustering patterns, but their expression trends appeared to diverge between sexes – showing opposite directions in males and females. Furthermore, the overall differences were modest and did not reach statistical significance in both sexes (Fig. 8g, h). For *Chil4* and *Wnt9a*, a trend toward decreased expression was seen in males but not in females. (Fig. 8f, i). Taken together, the combination of sex-dependent variability and the absence of significant differences in both sexes suggest that these genes are unlikely to be strong biomarker candidates.

**Fig. 8.**
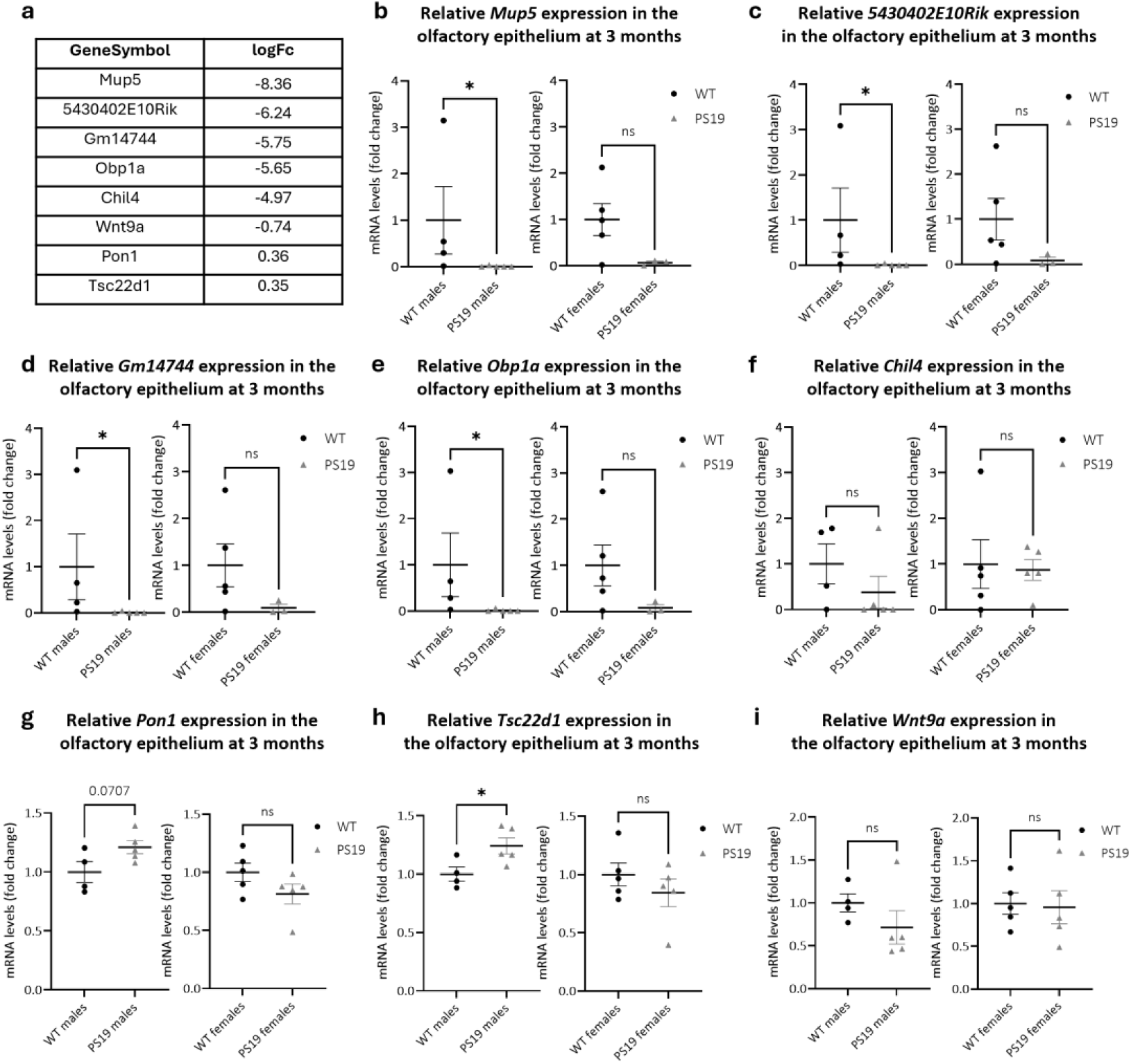
Biomarkers panel validation in the OE of 3-month-old PS19 and WT mice. Log fold-change (logFC) values were derived from RNA-sequencing analysis comparing PS19 mice to wild-type controls. Negative logFC values indicate downregulation, while positive values indicate upregulation in PS19 mice (a). mRNA expression levels of *Mup5*, *5430402E10Rik*, *Gm14744*, *Obp1a*, *Chil4*, *Pon1*, *Tsc22d1*, and *Wnt9a* were analyzed by RT-qPCR in the OE of WT and PS19 mice at 3 months of age (b-i). The results in males showed expression trends consistent with those observed in the RNA-sequencing data, although the differences did not always reach statistical significance (Unpaired t-tests, n = 3–5).

Next, we focused on *Gbp2b*, the only gene among the top 10 DEGs upregulated in the OE of 3-month-old PS19 males (Table 1, Fig. 9a). *Gbp2b* also ranked among the top 10 DEGs in the OE at 9 months, in the OB at 3 and 6 months, and in the hippocampus at 9 months (Table 1). Notably, we observed a highly significant increase in *Gbp2b* expression in the OE of both male and female PS19 mice at 3 months (Fig. 9b). In the OB, expression of *Gbp2b* was elevated as early as 3 months in both sexes and remained high at 9 months (Fig. 9c, d). In the hippocampus, we confirmed increased expression at 3 months in males and females, with the upregulation persisting at 9 months (Fig. 9e, f). So far, these findings suggest that this candidate represents the most promising early biomarker of tau pathology in the olfactory system.

**Fig. 9.**
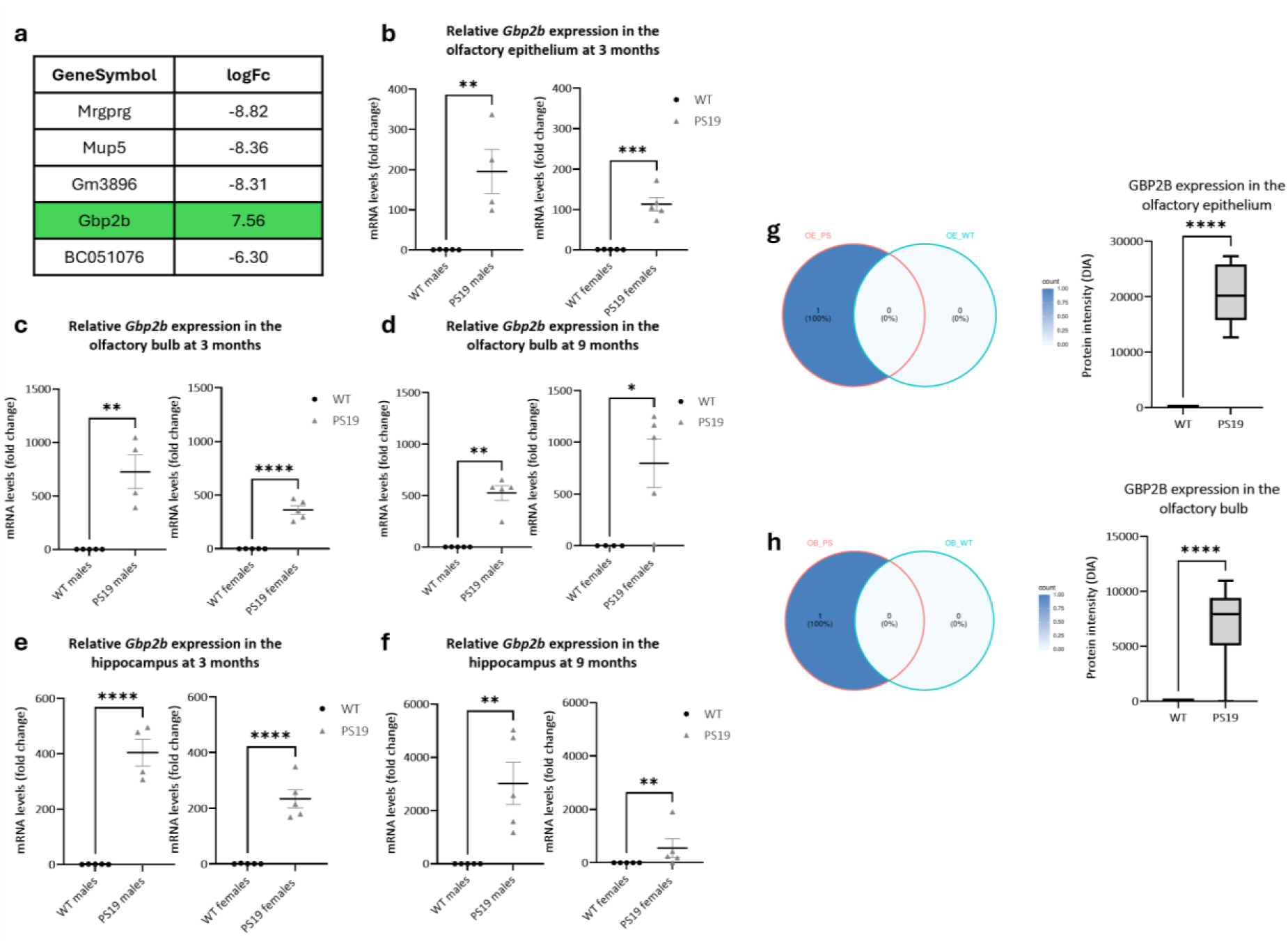
*Gbp2b* expression in the OE, OB and hippocampus of PS19 and WT mice. mRNA levels of *Gbp2b* expression were measured by RT-qPCR in OE, OB and hippocampal extracts from WT and PS19 mice at 3 months (for OE) and 9 months (for OB and hippocampus)(a-f). Top 5 DEGs in the OE of PS19 mice at 3 months with logFC values (a). *Gbp2b* expression is significantly increased in the OE of 3-month-old PS19 mice (b). \*\**P* < 0.01, \*\*\*\**P* < 0.0001 (Unpaired t-tests, n=4-5). *Gbp2b* expression is significantly increased in the OB of 3-month-old PS19 mice (c). \**P* < 0.05, \*\**P* < 0.01 (Unpaired t-tests, n=4-5). *Gbp2b* expression is significantly increased in the OB of 3-month-old PS19 mice (d). \**P* < 0.05, \*\**P* < 0.01 (Unpaired t-tests, n=4-5). *Gbp2b* expression is significantly increased in the HIP of 3-month-old PS19 mice (e). \*\*\*\**P* < 0.0001 (Unpaired t-tests n=4-5). *Gbp2b* expression is significantly increased in the HIP of 9-month-old PS19 mice (f). \*\**P* < 0.01 (Unpaired t-tests, n=4-5). **Box plot showing GBP2B protein intensity (log₂-transformed DIA signal) in the OE (g) and OB (h) of PS19 and WT mice.** Protein levels were quantified using data-independent acquisition (DIA) mass spectrometry. GBP2B was detected in all PS19 OE samples but was absent from WT samples. Boxes represent the interquartile range (IQR), horizontal lines indicate medians, and whiskers extend to 1.5× IQR. Statistical significance was assessed using an unpaired t-test (\*\*\*\**P* < 0.0001)(g). GBP2B was detected in 9 out of 10 PS19 OB samples but was absent from WT samples. Boxes represent the interquartile range (IQR), horizontal lines indicate medians, and whiskers extend to 1.5× IQR. Statistical significance was assessed using an unpaired t-test (\*\*\*\**P* < 0.0001)(h).

To further validate these transcript-level findings at the protein level, we opted for a data-independent acquisition (DIA) proteomic approach, which is known to outperform data-dependent acquisition (DDA) in the analysis of complex proteomic samples due to enhanced sensitivity for low-abundance peptides, increased specificity, and improved reproducibility [13].

Using DIA, we detected the mouse GBP2B protein in all OE samples from PS19 mice, while it was absent in all OE samples from WT mice (Fig. 9g). Similarly, GBP2B was detected in 9 out of 10 OB samples from PS19 mice and in none of the OB samples from WT mice (Fig. 9h). In the hippocampus, GBP2B was only detected in 3 samples out of 10 from PS19 mice, and absent in all WT mice samples (*data not shown*). These findings support the exclusive expression of GBP2B under PS19 condition and confirm that its upregulation is not limited to the transcriptomic level but is also reflected at the proteomic level.

Naturally, this targeted study evaluated only a limited subset of candidates. Additional potential biomarkers from the panel (Table 2) may warrant validation in future research.

## Discussion

Fifty million people worldwide currently suffer from NDs such as AD, and this number is expected to rise as life expectancy increases [27]. Despite extensive efforts to understand the underlying mechanisms and develop effective treatments, reliable early biomarkers for detecting these diseases before irreversible damage occurs remain elusive. Therefore, the primary goal of this study was to identify molecular signatures and biomarkers associated with tau pathology accumulation in the olfactory system, a uniquely accessible region with strong potential for monitoring CNS diseases.

The cerebral transcriptome of PS19 mice was previously established in the hippocampus [6] and cortex [15, 23] but data were still missing in regions of the olfactory system. Here, we used an integrated transcriptomic approach to map tau-driven molecular alterations in the OE, OB, and hippocampus at 3, 6, and 9 months of age.

### Region-Specific Molecular Changes Mirror Tau Pathology Progression in a Spatiotemporal Transcriptomic Analysis

Our results highlight a region-specific progression of tau pathology in PS19 mice, with distinct temporal and molecular signatures in the OE, OB, and hippocampus. In the OE, tau pathology is detectable as early as 3 months, persists at 6 months, and declines by 9 months, likely due to OSNs loss [11]. This trajectory corresponds with early transcriptomic changes involving stress responses, immune activation, and repression of olfactory transduction that persists over time. By 9 months, chronic inflammation, disrupted cilium organization, and metabolic disturbances dominate, suggesting that the OE’ regenerative capacity is overwhelmed by ongoing pathology [7]. Notably, chronic inflammation in this region may also arise as a downstream response to neuronal loss. These findings are consistent with transcriptomic profiles from human AD brains, which show downregulation of synaptic and neurotransmission-related genes alongside upregulation of immune responses and cell death pathways [24, 37].

In the OB, early changes affect sensory processing and neurogenesis, consistent with findings from transgenic AD models [38]. Although no major pathway enrichment is observed at 6 months, the 9-month time point reveals mitochondrial dysfunction, ER stress, and emerging sensory deficits. Pathologically, the OB remains relatively stable with a stall in pathological progression where tau remains in a hyperphosphorylated state without aggregating further. This limited progression may explain the delayed but modest transcriptional response, despite early evidence of synaptic dysfunction. It is even more striking knowing that soluble forms of pathological tau rather than aggregated tau are suggested to induce synaptic dysfunction [8, 39].

In contrast, the hippocampus shows a more classic, gradual accumulation of tau pathology from 3 to 9 months, reflected in a steady increase in DEGs. Early disruptions in synaptic plasticity and neuronal function evolve into widespread neurodegeneration and impaired neurotransmission by late stages, aligning with established role of the hippocampus in cognitive decline and with previous observations in PS19 mice [6].

### Early and Region-Specific Gene Expression Changes in PS19 Mice Highlight Potential Biomarkers for Tauopathies

Overlap analysis of DEGs across regions and time points revealed both shared and region-specific responses. The earliest and most distinct molecular changes were observed in the OE—the first region to show tau pathology. We focused on this region at 3 months of age and identified several DEGs, including *Gbp2b, Prnp, Wnt9a, Mup5, Chil4,* and *Obp1a* (Table 1). Some, like *Wnt9a* and *Prnp*, have been previously associated with AD or reported in other tauopathy models [1, 6]. *Wnt9a*, part of the Wnt signaling pathway critical for synaptic plasticity [25], was upregulated in the hippocampus and OB but downregulated in the OE. Similar enrichment of synaptic, mitochondrial, and calcium-related genes in the hippocampus parallels prior findings in the hippocampus and cortex of PS19 mice [1]. In the cerebral cortex of PS19 mice, overexpression of aggregation-prone tau alone has been reported to trigger early neuroinflammatory responses—including upregulation of immune-related genes like *Aqp1, Lbp, Prnp,* and *Hsb1*—as early as 3 months [15]. We observed similar changes, alongside upregulation of apoptotic and mitochondrial genes, reinforcing the idea that tau pathology initiates broad, system-wide immune and metabolic responses.

Among these DEGs in the OE, *Gbp2b* emerged as a particularly compelling biomarker due to its consistent transcript-protein match, temporal persistence, and cross-region upregulation. This is supported by earlier reports showing elevated GBP2 – belonging to the GBP family – expression in the cortex, hippocampus, and OB of 5xFAD mice and its involvement in microglial activation and neuronal apoptosis after brain injury [21, 23]. As an interferon-induced gene, GBP2 is known to be activated early in the inflammatory response and has been proposed as a biomarker and therapeutic target in AD models [42]. The early vulnerability of olfactory regions to tau pathology may explain the strong and consistent upregulation of *Gbp2b* observed in these areas. This candidate is particularly striking given that mouse Gbp2b and human GBP1/GBP2 proteins are true orthologs, sharing a high degree of sequence similarity—approximately 70%—which underscores their likely functional conservation.

We found that genes downregulated in the OE at 3 months (e.g., *Mup5, Obp1a, 5430402E10Rik,* and *Gm14744*) are linked to odorant binding and chemical communication — processes critical for olfactory function. We also observed reduced expression of *Chil4*, which is involved in OE regeneration via inflammatory pathways [36]. Impairments in these biological processes may contribute to the progressive olfactory deficits often observed in AD patients [2, 3, 44] and supports a link between tau pathology and olfactory dysfunction. Internal validation of these candidates confirmed a decrease expression in the OE at 3 months in both PS19 males and females.

### Nasal Brushing as a Minimally Invasive Strategy for Early Detection of Tauopathy Biomarkers

To date, biomarkers are collected through invasive techniques, especially CSF obtained through lumbar puncture [26]. Blood-based biomarkers (BBMs) were also proposed for diagnosing AD, but they are not yet accurate enough to replace CSF and PET testing [5]. In recent years, nasal brushing has garnered significant interest as a minimally invasive and straightforward technique for assessing OE-derived biomarkers in NDs [12, 17, 29]. In the search for reliable biomarkers, those that are upregulated under pathological conditions — such as *Gbp2b* — are generally preferred. Indeed, elevated expression levels are easier to detect and interpret, offering greater analytical sensitivity [34]. In contrast, biomarkers with decreased expression can be more challenging to measure accurately, increasing the risk of false negatives due to low signal levels [14, 35].

A promising avenue is to validate the biomarkers identified in our tauopathy mouse model by determining whether these markers can be detected in nasal brushing samples and effectively discriminate between control subjects and early-stage AD patients. Given that the OE is one of the first regions affected by AD pathology and maintains a direct connection to the CNS [31, 33], OE-derived biomarkers may offer earlier and more reliable diagnosis of NDs.

In conclusion, our integrated transcriptomic profiling reveals that tau overexpression and aberrant phosphorylation in PS19 mice leads to early, region-specific molecular changes—particularly in the OE. These changes begin with olfactory dysfunction, inflammation, and cellular stress at 3 months, and progress to broader disruptions in neuronal signaling and metabolism by 9 months. We identified several potential biomarkers, with *Gbp2b* standing out due to its consistent upregulation, ease of detection, and established relevance in inflammation and neurodegeneration. Importantly, *Gbp2b* expression was validated in both sexes and across multiple time points and brain regions. These findings underscore the diagnostic potential of OE-derived biomarkers and highlight nasal brushing as a promising and viable method for early detection. Ultimately, this approach may accelerate the development of predictive diagnostics and targeted interventions for tau-related NDs.

#### Limitations of this study

We used a number ranging from 3 to 5 males per condition for transcriptomic studies. Increasing the sample size could improve the robustness and accuracy of the data. Still, further validation (proteomic and RT-qPCR) indicated that to-differentially-expressed candidates were accurately identified in the transcriptomic signatures.

Given that the identified potential biomarker genes participate in diverse biological pathways, once validated (e.g. in nasal brushing samples), it is likely that a panel comprising multiple genes would serve as a more robust diagnostic tool than any single candidate.

## Declarations

### Statement on welfare

All animal procedures were conducted in accordance with institutional and European guidelines and approved by the UCLouvain Ethical Committee for Animal Welfare (2021/UCL/MD/018).

### Data available statements

All datasets generated and analyzed during this study are included in this published article and its supplementary material. RNA-sequencing data have been submitted to SRA and will be made publicly available upon acceptance. Proteomics data have been submitted to PRIDE and will be made publicly available upon acceptance. Accession numbers will be provided upon publication.

### Authors contribution

Conceptualization: MD, AB, PR, PKC; Methodology: MD, AB, EP, MM, MD; Formal analysis and investigation: MD, AB, MM, MD; Writing – original draft preparation: MD, AB; Writing – review and editing: MD, AB, MD, NS, PR, PKC; Funding: PKC; Supervision: PR, PKC. All authors read and approved the final manuscript.

## List of Abbreviations

ABs: Axon bundles
AD: Alzheimer’s disease
CNS: Central nervous system
DEGs: Differentially expressed genes
NDs: Neurodegenerative diseases
OB: Olfactory bulb
OE: Olfactory epithelium
OSNs: Olfactory sensory neurons
WT: Wild-type

