## Supplementary information for "An Early Olfactory Transcriptomic Signature of Tauopathy: Gbp2b Emerges as a Candidate Biomarker of Tau-Driven Neuroinflammation"

*Contributed equally

**Supplementary figures and tables**

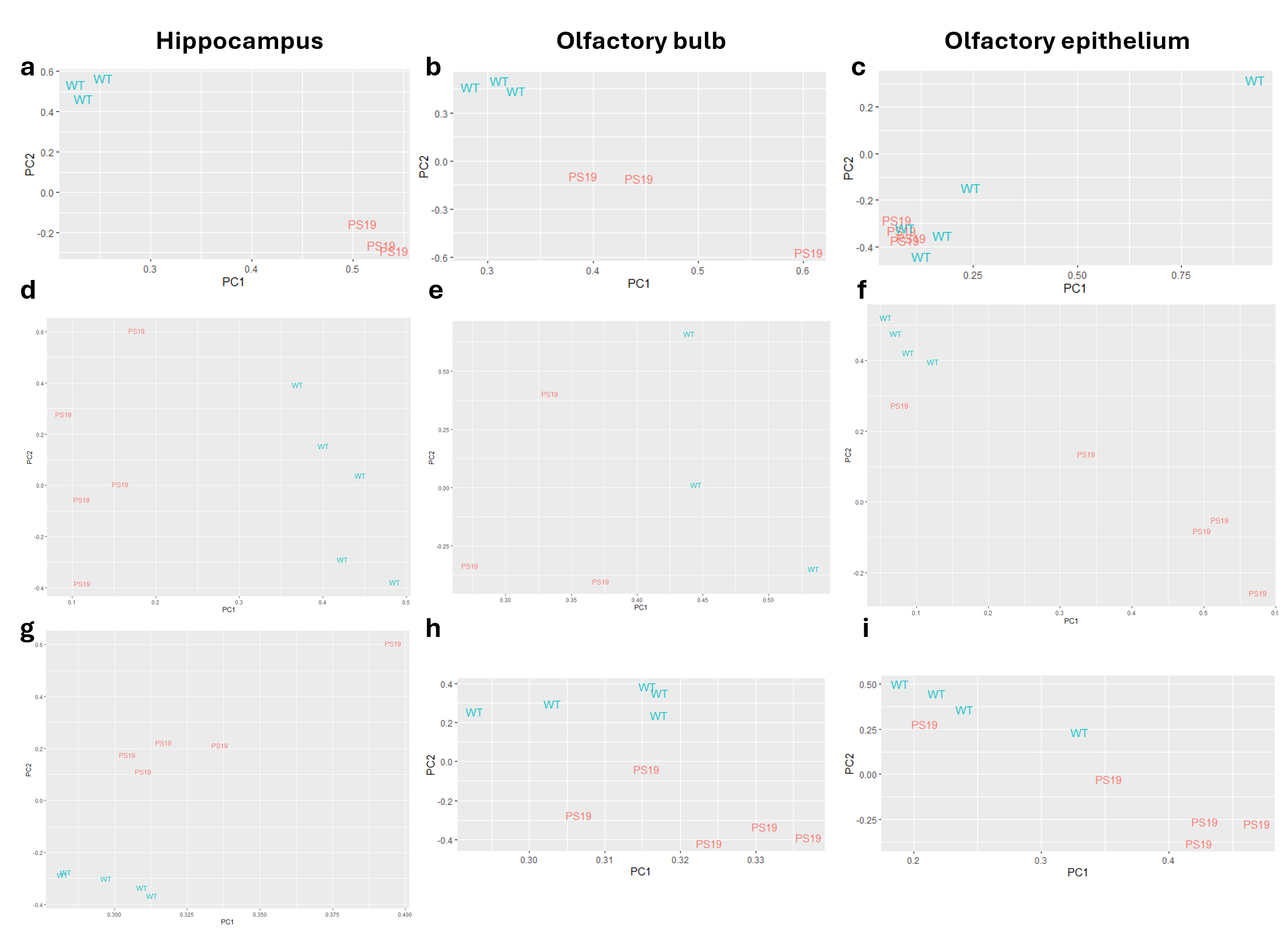
 **Fig. S1 PCA of normalized gene expression in hippocampus (a, d, g), OB (b, e, h), and OE (c, f, i) from PS19 and WT mice at 3 (a-c), 6 (d-f), and 9 (g-i) months.** Each point represents a sample, colored by genotype (PS19 in pink, WT in blue). The first two principal components (PC1 and PC2) are shown.

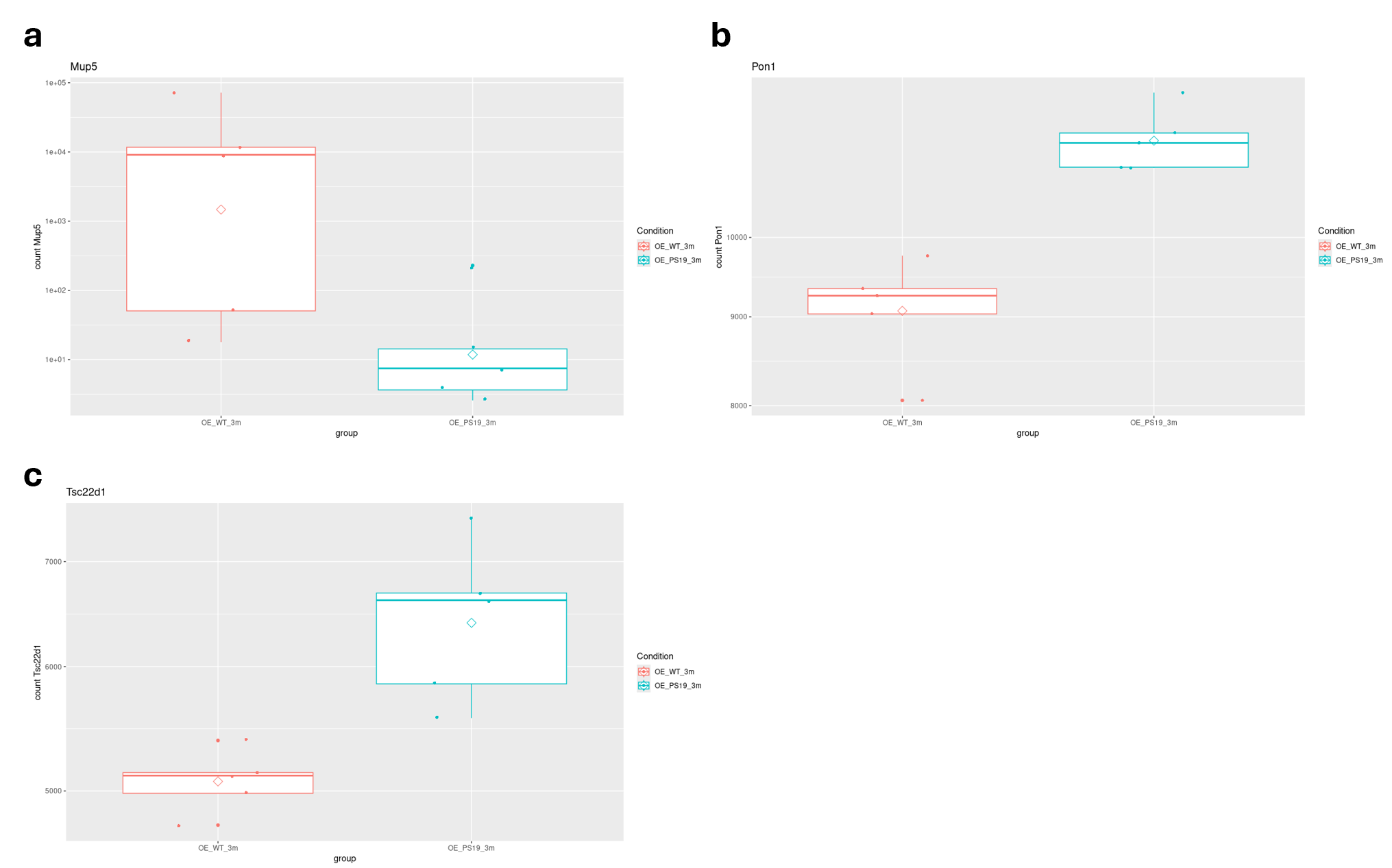
 **Fig. S2 Expression levels of Mup5, Pon1, and Tsc22d1 in the OE of WT and PS19 mice.** Boxplots show normalized expression counts for three genes—Mup5, Pon1, and Tsc22d1—in two experimental groups: OE_WT_3m (red) and OE_Ps19_3m (cyan). Each plot displays the distribution of gene expression levels across biological replicates. Boxes represent the interquartile range (IQR), with horizontal lines indicating the median and diamonds showing the mean. Whiskers extend to 1.5 times the IQR, and individual data points are overlaid to illustrate sample variation.

| **Gene** | **Forward Primer** | **Reverse Primer** |
| --- | --- | --- |
| Gapdh | 5'-ACCCAGAAGACTGTGGATGG-3' | 5'-ACACATTGGGGGTAGGAAC-3' |
| Prnp | 5’-CCCGCGTTGTCGGATCA-3’ | 5’-AGAATGCTTCAGCTCGGTTG-3’ |
| Mup5 | 5’-TCCAGTGTTGAGTGGAGACT-3’ | 5’-CTTGTCACCCCATGCTGTAT-3’ |
| 5430402E10Rik | 5’-GCGCCCTTTTCTTTCACAAC-3’ | 5’-TTGCTCAGGGTCTCTCCTTT-3’ |
| Gm14744 | 5’-CGCCCTTTTCTTTCACAACG-3’ | 5’-TGCTCAGGGTCTCTCCTTTT-3’ |
| Obp1a | 5’-AGCTGGAAACATCAGAGAAGTC-3’ | 5’-GTCAAGACCACATGATCATAATTC-3 |
| Chil4 | 5’-GCCAAGCTCATTCTTGTCAC-3’ | 5’-ACACAGGCAGGGGTCAATA-3’ |
| Pon1 | 5’-GGACTGGTGTTGGCACTTTA-3’ | 5’-CTGGCGTTACTTCACGGAAA-3’ |
| Tsc22d1 | 5’-CTCTGGTGCAAGTGTGGTAG-3’ | 5’-CCCTCACCGCATACATCAAA-3’ |
| Wnt9a | 5’-GTGGACTTCCACAACAACCT-3’ | 5’-TGGCATTTGCAAGTGGTTTC-3’ |
| Gbp2b | 5’-AGACTCCTGGAAAGGGACTC-3’ | 5’-TGTGAGTGACTGGTCCATCT-3’ |

**Table S1. qPCR primer sequences.**
